# The action of Con-ikot-ikot toxin on single AMPA-type glutamate receptors

**DOI:** 10.1101/2021.02.19.432041

**Authors:** Jelena Baranovic, Sebastian Braunbeck, Nikolai Zaki, Sonja Minniberger, Miriam Chebli, Andrew J.R. Plested

## Abstract

Conotoxins are a large group of naturally occurring toxic peptides produced by the predatory sea snails of the genus *Conus.* Many of these toxins target ion channels, often with high specificity and affinity. As such, they have proven to be invaluable for basic research as well as acting as leads for therapeutic strategies. Con-ikot-ikot is the only conotoxin so far identified that targets AMPA-type glutamate receptors, the main mediators of excitatory neurotransmission in the vertebrate brain. Here, we describe how the toxin modifies the activity of AMPA receptors at the single-channel level. The toxin binds to the AMPA receptor with high affinity (*EC*_50_ = 5 nM) and once bound, takes minutes to wash out. As shown previously, it effectively blocks desensitization of AMPA receptors, however, compared to other desensitisation blockers, it is a poor stabiliser of the open channel because toxin-bound AMPA receptors undergo frequent, brief closures. We propose this is a direct consequence of the toxin’s unique binding mode to the ligand binding domains. Unlike other blockers of desensitization, which stabilise individual dimers within an AMPA receptor tetramer, the toxin immobilizes all four ligand binding domains of the tetramer. This result further emphasises that quaternary reorganization of independent LBD dimers is essential for the full activity of AMPA receptors.

## Introduction

Predatory sea snails of the genus *Conus* are a rich source of toxic peptides known as conotoxins. Conotoxins are mostly disulphide-rich peptides which target invertebrate and vertebrate ion channels with high specificity. With >500 species of *Conus* snails and each snail producing a venom composed of >100 different peptides, there are more than 50 000 different, pharmacologically-active conotoxins (Terlau & Olivera, 2004). This vast natural pharmacy has proven vital for basic research on ion channels (Safo et al., 2000; Xiong et al., 2020), as well as for the development of various therapeutics (Essack et al., 2012). Ziconotide is the first FDA approved drug based on a conotoxin and is used in the treatment of severe chronic pain. A number of conotoxin-based drugs are currently in clinical trials for various pathologies, such as a peptide based on a conantokin G, a conotoxin that targets NMDA receptors, for the treatment of epileptic seizures (Essack et al., 2012). Majority of conotoxins target voltage and ligand gated ion channels, with acetylcholine receptors the most common target among the latter.

The discovery of Con-ikot-ikot toxin (CII) added the AMPA (a-amino-3-hydroxyl-5-methyl-4-isoxazole-propionic acid) type glutamate receptors (AMPA receptors) to the repertoire of conotoxin targets (Walker et al., 2009). Con-ikot-ikot (from C. *striatus)* is an unusual member of the conotoxin superfamily. Although it is rich in the typical disulphide linkages, their pattern is distinct from that in any existing conotoxin family and it contains 86 vs. usual 20-30 amino acids (Fig. 1A-B) (Robinson & Norton, 2014).

**Figure 1.**
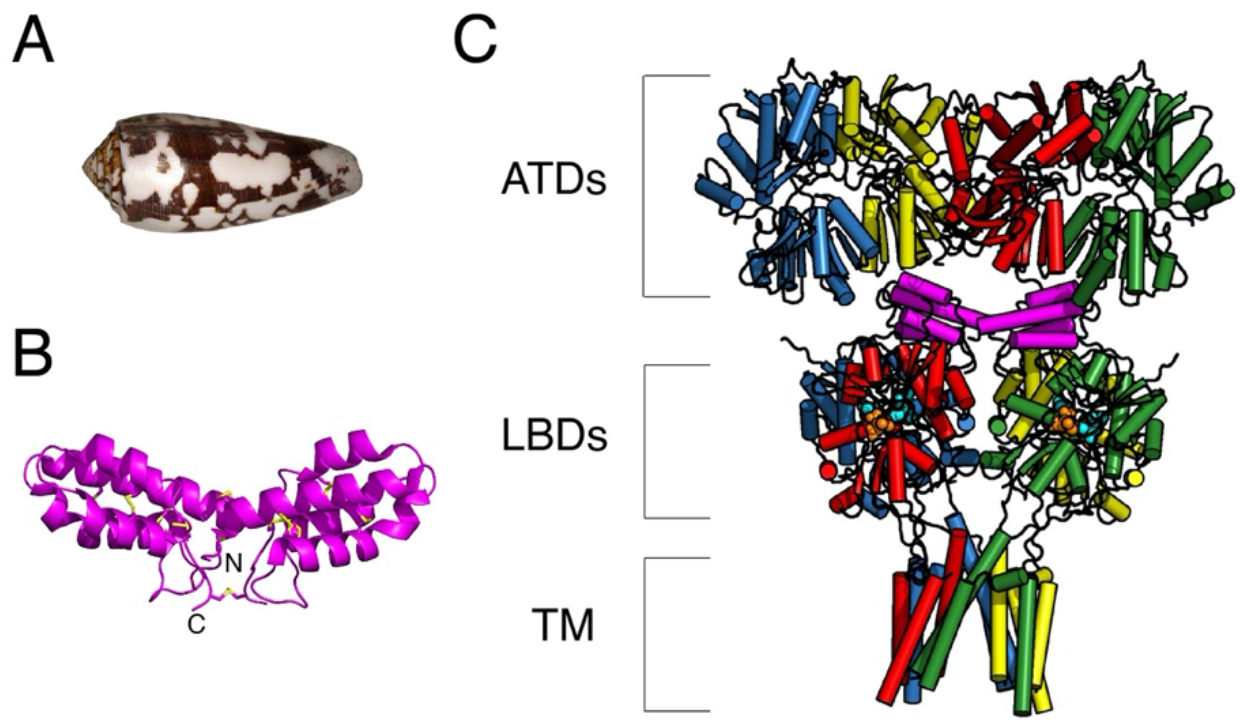
Structure of Con-ikot-ikot toxin. (A) Shell of a *Conus striatus* snail, the natural source of Con-ikot-ikot toxin (CII). (B) Crystal structure of the toxin homodimer; each CII monomer is a 4 helix bundle containing 5 disulphide bridges, connected to another monomer via additional 3 disulphide bridges (yellow) (PDB: 4U5H (Chen et al., 2014)). (C) Crystal structure of the full-length AMPA receptor (subunit GluA2) in complex with a toxin (magenta), partial agonist kainate and desensitization blocker (R, R)-2b (PDB: 4U5D (Chen et al., 2014)); each AMPA receptor subunit is coloured differently with the domains indicated with square brackets as: ATDs – amino terminal domains, LBDs – ligand binding domains and TM – transmembrane region. Red and blue subunits are forming one and green and yellow the other LBD dimer. The V-shaped toxin dimer (magenta) sits on top of LBDs.

AMPA receptors are the main mediators of excitatory neurotransmission in the vertebrate central nervous system. Their fast kinetics generally make them the first glutamate receptor subtype in the postsynaptic membrane to be activated by glutamate released from synaptic vesicles. Once bound by glutamate, their integral ion channel opens to allow Na^+^ influx, initiating membrane depolarisation. Binding of glutamate, however, keeps the channel open for only a fraction of a millisecond because the receptor inactivates rapidly despite glutamate still being bound, a phenomenon known as desensitization. Desensitization plays roles in normal synaptic transmission (DiGregorio et al., 2007; Jones & Westbrook, 1996) and appears important for nervous system development (Christie et al., 2010).

AMPA receptors are tetrameric proteins composed of four different subunits, GluA1-A4. Each subunit is composed of amino terminal domain (ATD), ligand binding domain (LBD), transmembrane segment with the ion channel and carboxy terminal (Fig. 1C) (Sobolevsky et al., 2009). ATDs and LBDs form the extracellular part of the receptor, protruding into the synaptic cleft. ATDs are separated from LBDs by flexible linkers, leading to practically no interaction between the two domains and creating free space between them (Yelshanskaya et al., 2016). As shown by the crystal structure of an AMPA receptor in complex with the Con-ikot-ikot toxin, this cavity between ATDs and LBDs in the tetramer is exactly where the toxin dimer binds, interacting mainly with the LBDs (Fig. 1C) (Chen et al., 2014). The V-shaped toxin dimer fits neatly on top of the AMPA receptor LBDs: four LBDs in the receptor are arranged as a dimer-of-dimers and each toxin monomer sits on top of one dimer. Thus, as long as the toxin is bound, the dimers remain intact. This is important, as in order to desensitize, the receptor needs to break the interface within individual dimers (Sun et al., 2002). As a consequence, CII toxin blocks AMPA receptor desensitization, prolonging its activation and over-exciting the postsynaptic neuron (Walker et al., 2009). This mechanism is the basis of CII toxicity. Given the block of desensitization, it was somewhat surprising to see that crystal structures of an AMPA receptor in complex with CII toxin, partial agonist and another desensitization blocker resulted in a closed ion channel pore (Chen et al., 2014).

Here, we explore the mechanism of CII toxin on AMPA receptor activity at the single-channel level. The toxin binds to AMPA receptors with nM affinity and blocks desensitization effectively. However, when compared to other desensitization blockers, such as cyclothiazide (CTZ) and (R, R)-2b, the toxin is a poor stabiliser of the open channel, leading to frequent and brief closures. We propose the reason for this inability to stabilise the active state of the receptor lies in the toxin’s unique mode of binding, distinct from other desensitization blockers. Whereas CTZ and (R, R)-2b both bind within individual dimers, leaving them free to move with respect to each other, the toxin “locks” the two dimers at a fixed angle. Thus, the toxin fully immobilizes the LBD layer of the receptor, which appears incompatible with a stably open ion channel.

## Materials and Methods

### Toxin Expression and Purification

The plasmid pET32b containing the mature sequence for the cone snail toxin Con-ikot-ikot (amino acids 38-123, UniprotKB number P0CB20), preceded by a Trx fusion tag, a Strep tag and a HRV 3C cleavage site was a kind gift of Eric Gouaux (Chen et al., 2014). The vector was transformed into Origami B (DE3) *E. coli* cells and grown in LB medium until OD600 reached 0.8 – 1.0. The cultures were then induced with 100 μM IPTG at 16°C and harvested 16-20h post-induction. For cell lysis, the pellet was resuspended in lysis buffer (50 mM Tris-HCl pH 8, 150 mM NaCl, 5 mM EDTA, 1 mM Pefablock, 50 μg/ml lysozyme and 25 μg/ml DNaseI) and sonicated on ice for 5 min in total. To collect the supernatant, the lysate was centrifuged at 20 000 rpm with a Fiberlite F21-8×50y rotor for 40 min and loaded onto a YMC ECO15/120V0V column packed with 10 ml Strep-Tactin Superflow high capacity resin. First, the column was washed extensively with Strep buffer A (20 mM Tris-HCl pH 8, 150 mM NaCl, 1 mM EDTA) and then eluted in one step with Strep-buffer B (20 mM Tris-HCl pH 8, 150 mM NaCl, 1 mM EDTA, 2.5 mM *d*-desthiobiotin). A 30 kDa cut-off Amicon centrifugal filter was used to concentrate the eluted protein and 3C protease (1:100) added to cleave off the Trx tag overnight at 4°C. The next day, 2 volumes of methanol were added to the digested protein in order to precipitate the Trx tag, incubated at 37°C for 10 min and centrifuged at 4 000 rpm for 10 min. The supernatant was recovered and concentrated using a 3 kDa cut-off Amicon centrifugal filter, exchanging the buffer with consecutive concentration runs to 20 mM Tris-HCl pH 8, 150 mM NaCl, 1 mM EDTA. The resulting concentrate was left at room temperature for 24h before addition of 1 mM GSH and further incubation at room temperature for another 24h to promote dimer formation. The GSH treated sample was diluted 10-fold with SP_A buffer (30 mM NaAc pH 4.2) and loaded onto a HiTrap SP HP 1 ml prepacked column. The protein was eluted using a gradient from 100 mM to 250 mM NaCl. Fractions of the ion exchange run were submitted to SDS-PAGE to identify fractions containing functional toxin dimers. These fractions were concentrated using a 3 kDa cut-off Amicon centrifugal filter and loaded onto a Superdex 75 10/300 column equilibrated in 10 mM HEPES pH 7.5, 150 mM NaCl (Fig. 2A). Toxin dimer was identified by SDS-PAGE (Fig. 2B) and the corresponding fractions were pooled and concentrated using a 3 kDa cut-off Amicon centrifugal filter to a final concentration of 0.9 mg/ml. The toxin was stored at 4°C after the addition of 0.01% NaN3 until further use.

**Figure 2.**
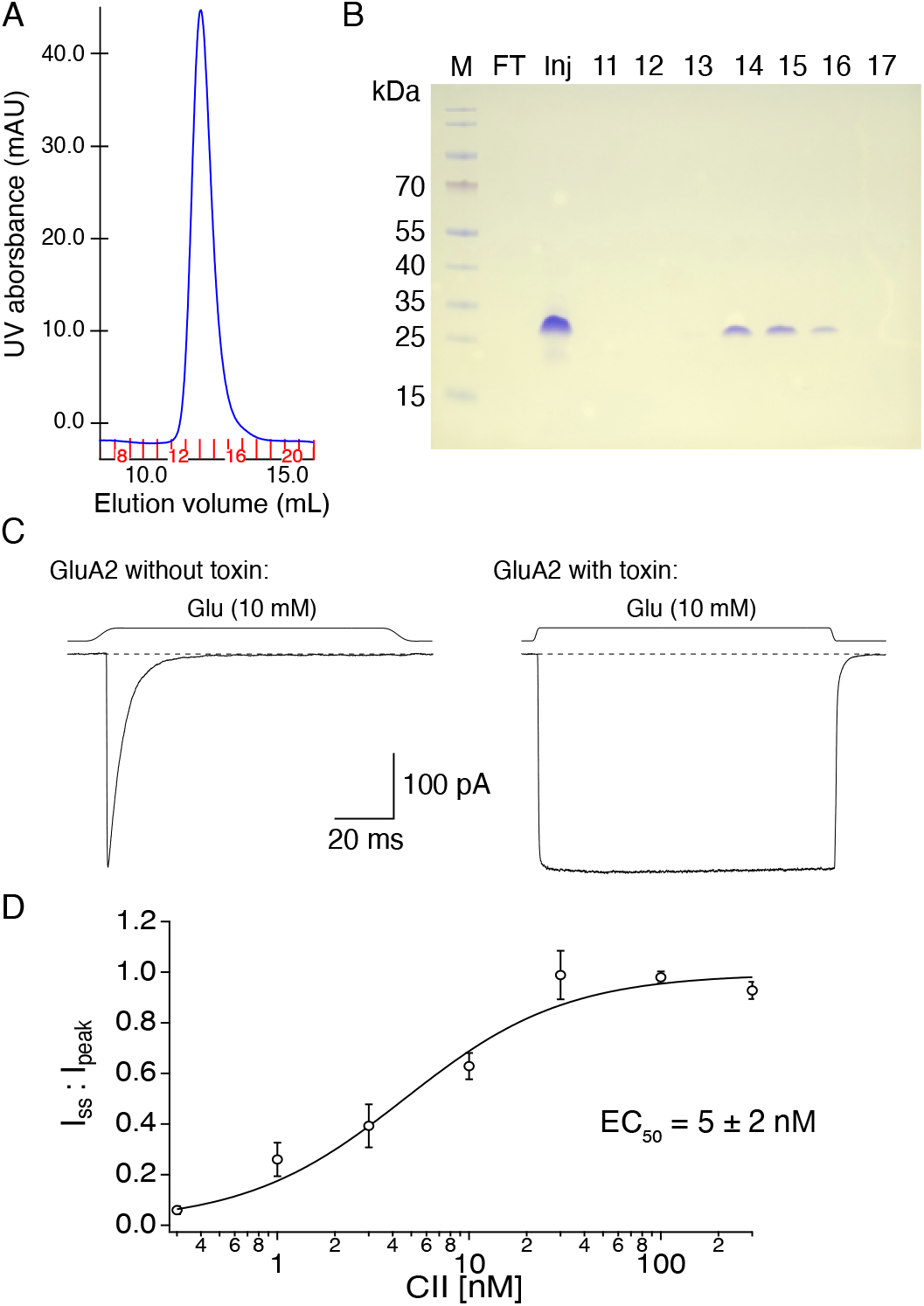
Con-ikot-ikot toxin purification and characterization. (A) Size-exclusion chromatogram (SEC) of purified Con-ikot-ikot toxin (CII) with fraction numbers indicated in red. (B) Coomassie-stained non-reducing SDS-PAGE gel of the SEC fractions. Wells marked 11-17 are SEC fractions, “Inj” is the concentrated sample injected onto the SEC column and FT is the flow-through after concentrating the sample and before loading it onto the SEC column (to check for any protein loss). Fractions 14-16 contain purified CII dimer. (C) Activity of the purified toxin was checked with outside-out patches: left trace shows current produced by GluA2 AMPA receptors in the presence of glutamate (10 mM) without CII bound and right trace is from a different patch with CII toxin bound (after incubation in 100 nM toxin). Dashed line indicates baseline and square trace above it solution exchange in the respective patch. Voltage was ~ —40 mV in both recordings. (D) CII toxin dose response curve for AMPA receptors (0.3-300 nM CII, *n* = 3-8 patches/toxin concentration). Toxin effect was measured as desensitization block, i.e. ratio of steady-state current over peak current for each patch (1 for complete desensitization block). Fit by Hill equation gave *EC*_50_ of 5 ± 2 nM and Hill slope of 1 ± 0.3 (95% confidence interval).

### Electrophysiology

All recordings were performed with the wild-type GluA2 subunit of AMPA receptors in flip form, unedited (Q) at the Q/R site in the channel pore. EGFP was present downstream from the GluA2 coding region after an IRES to mark transfected cells. Wild-type AMPA receptors were expressed transiently in HEK293 cells using calcium-phosphate precipitation or PEI method as described previously (Baranovic et al., 2016; Riva et al., 2017). For macroscopic recordings, cells were recorded 48-72 hours post transfection and for single-channel recordings, 24 hours post transfection. In both cases we used the outside-out patch configuration.

Recording pipettes were pulled from borosilicate glass and had resistances (when filled with internal solution) of 5-10 MΩ for macroscopic recordings and 10-20 MΩ for single-channel recordings. The internal (pipette) solution in all cases contained (in mM): 115 NaCl, 1 MgCl_2_, 0.5 CaCl_2_, 10 NaF, 5 Na_4_BAPTA, 10 Na_2_ATP and 5 HEPES, titrated to pH 7.3 with NaOH. The external solution consisted of (mM): 150 NaCl, 0.1 MgCl_2_, 0.1 CaCl_2_ and 5 HEPES, titrated to pH 7.3 with NaOH. Desensitization blockers CTZ and (R, R)-2b were added to the external solution at the final concentration of 100 and 10 μM, respectively, where noted. Stock solutions of both drugs were prepared in DMSO (at 50 mM and 100 mM, respectively) and stored at – 20°C. CTZ was obtained from Hello Bio and (R, R)-2b was a kind gift from Eric Gouaux. Receptors were activated with glutamate (10 mM) solution that also contained 5 mM sucrose to visualize the flowing stream from the fast perfusion tool.

Con-ikot-ikot toxin was either present in the perfusing external solution at 500 nM when observing binding of the toxin or in the bath external solution at saturating concentrations (350 nM – 45 μM) for measuring the unbinding rate and performing trace idealisation, with or without additional desensitization blockers. For constructing the dose response curve (Fig. 2D), HEK293 cells expressing wild-type GluA2 receptors were incubated in culture dishes overnight at various toxin concentrations.

We applied glutamate (with and without toxin, CTZ or (R, R)-2b) to outside patches via perfusion tools made from custom-manufactured four-barrel glass (Vitrocom) (Plested & Mayer, 2009). The 10%-90% solution exchange time was measured from junction potentials at the open tip of the patch pipette at the end of each experiment and was <300 μs. Patches were clamped at –40 to –60 mV for macroscopic records and at –80 mV for single-channel currents, unless stated otherwise. For single channel recordings, we regularly applied gentle suction to deform the patch and reduce the number of active channels. Currents were filtered at 10 kHz (–3 dB cutoff, 8-pole Bessel, Axopatch 200B amplifier) and recorded using AxoGraph X (Axograph Scientific, v1.7.6) via an Instrutech ITC-18 interface (HEKA) at 20 kHz sampling rate.

### Analysis of macroscopic recordings

Ad hoc analysis during experiments and pre-processing of data was done in AxoGraph. Traces were averaged, baseline-corrected and selected for fitting of exponentials before export. For the toxin dose response curve (Fig. 2D), the ratio of steady-state current to the peak current (I_SS_ / I_peak_) was determined for each patch at a range of toxin concentrations (0.3-300 nM, *n* = 3-8 patches per toxin concentration). Data were fit in Igor Pro (v7.08) with a Hill equation:

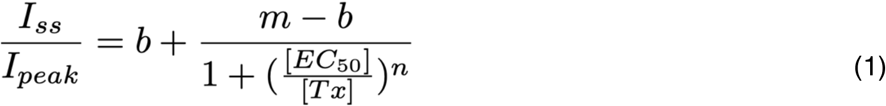

where *b* is the baseline steady-state current without toxin (fixed to 0.01 for fit), *m* is the maximum block of desensitization, [*EC*_50_] is the midpoint concentration, *n* is the Hill slope and [*Tx*] is toxin concentration.

### Single-channel recordings

For single-channel analysis, records were digital Gaussian filtered and selected for export with AxoGraph. The time needed for Con-ikot-ikot toxin to unbind from single GluA2 wild-type receptors was determined by perfusing toxin-bound receptors in a toxin-free external solution (containing glutamate only) until toxin unbinding was observed as a sharp drop in the number of channel openings (Fig. 3B).

**Figure 3.**
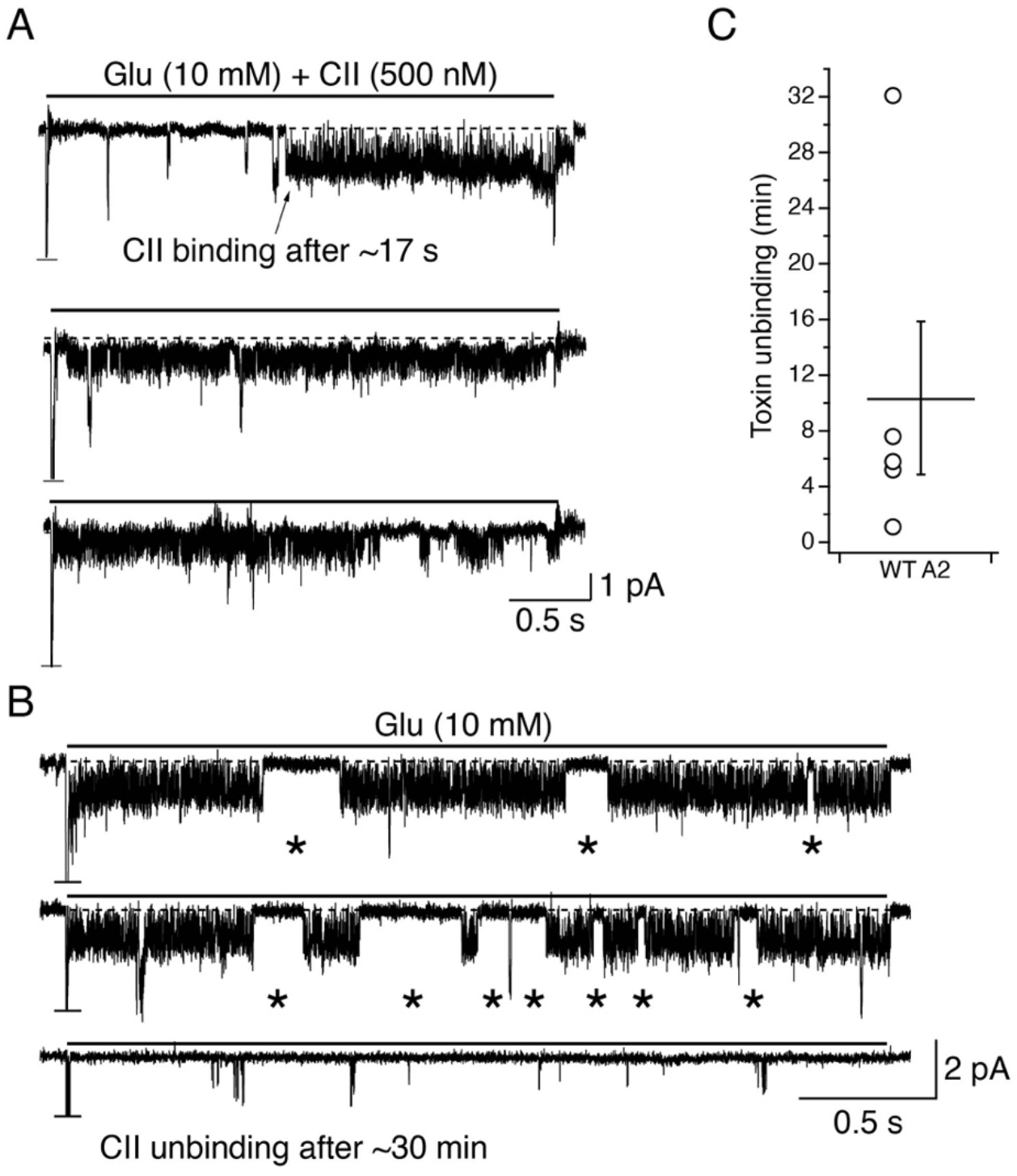
Binding and unbinding of toxin to individual AMPA receptors. (A) Current trace showing binding of toxin to one AMPA receptor in a patch containing multiple receptors, as indicated by the truncated peak current response at the start of glutamate application (horizontal line). Arrow indicates toxin binding event after 17 s. (B) To detect toxin unbinding, patches with bound toxin were perfused in a toxin-free solution. In this example, long bursts of activity characteristic for toxin-bound AMPA receptors (top two traces) were replaced by short bursts of activity interspersed by long closures (bottom trace), after about 30 min. In both, A and B, dashed line indicates baseline and solid, black line above the traces, length of glutamate application. Voltage was clamped at –80 mV for both traces. Asterisks mark longer shut epochs (>23 ms) during the toxin bound phase. (C) After binding of toxin to one AMPA receptor in a patch, it took 10.4 ± 5.5 min (*n* = 5) to observe unbinding in a toxin-free solution, with regular 3 s jumps into glutamate.

Amplitude points from the duration of agonist application (3 s) were concatenated into a single (continuous) current trace. An all-point-amplitude histogram was then generated for every patch in Igor Pro (v7.08), fit with a multi-peak Gaussian mixture function and normalised. Fits from various patches belonging to the same condition (Glu + CTZ, Glu + CII, Glu + (R, R)-2b and Glu + CII + (R, R)-2b) were plotted on the same graph to show variability across patches within a condition and to compare the four conditions.

Idealisation of single-channel records was performed only on patches with baseline root mean square noise of less than 300 fA, in a PYTHON-based, open source single-channel analysis application with a graphical user interface (ASCAM, www.github.com/AGPlested/ASCAM). ASCAM was optimised to handle multi-episode files and allow the user to select and mark traces for further analysis. Traces were saved as .mat files (the MATLAB native format, for compatibility with other third-party software) and imported back into AxoGraph or Igor Pro for further analysis. Traces were filtered and the baseline was subtracted using a linear fit to regions of baseline. Detection of open and closed channel events was limited to the interval over which agonist was applied by automatically thresholding the piezo command voltage. This procedure excluded any spurious detection from intervals of baseline. Due to desensitization block, in most patches, the channel was open as long as the agonist was present; however, occasionally, in 3/7 patches, shuttings longer than 23 ms were observed, which were excluded from idealisation. Thus, the longest intra-burst shut time was 23 ms. We regularly observed 4 open levels for each active AMPA receptor. Idealisation of single-channel records was done in ASCAM with a multiple threshold-crossing algorithm after filtering and baseline-correction of the current traces. Each threshold was taken from the bisector of the adjacent open levels, and the bisector of the first open level and the zero baseline shut level. These open levels were specified by the user from inspection of the all-point histograms. Dead time (duration of the shortest event that could be accurately detected) during idealisation was set to 130 μs. We typically used interpolation (5-fold) to improve the determination of the threshold crossing time. The idealisation process returned a numbered list of events (of channel opening times, levels and the interspersed closed times). The list of events for each patch was exported as text-file for further analysis in Igor Pro.

Dwell time intervals (*t*) were log-transformed (equation 2) (Sigworth & Sine, 1987),

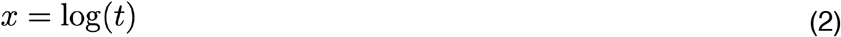

The probability density function was appropriately transformed (equation 3),

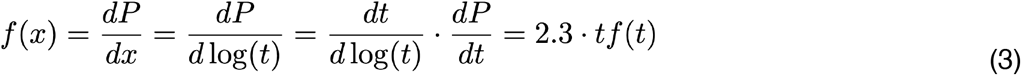

Dwell time histograms were then fitted with equation 4.

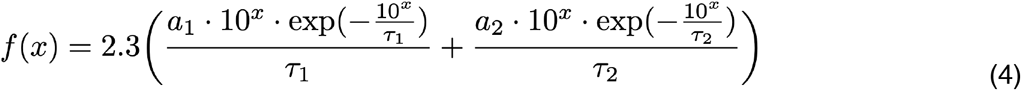

where *τ_1_* and *τ_2_* are time constants of the exponential components of the fit.

To determine “fractional activity” of a single GluA2 receptor in complex with different desensitization blockers, we calculated fraction of the maximum charge (*Q*_Frac_) transferred during the single-channel activity using equation 5:

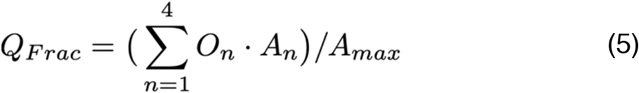

Where *O_n_* is the fraction of time spent at each open level *n* (1-4), determined from the idealization, and *A*_n_ is the amplitude of that level; *A*_max_ is the amplitude of the maximum open level in the recording (that is, *A*_4_). Maximum charge is defined as the charge that would be transferred across the channel if the channel was open at *A*_max_ for the whole duration of agonist application.

All *P* values were determined by a non-parametric randomization test (using ≥ 10^5^ iterations; DCStats suite, https://github.com/aplested/DC-Stats) The spread of the data is indicated as standard deviation of the mean unless stated otherwise. Graphs were plotted and fitted in Igor Pro (Wavemetrics). Visualisation of molecular structures was done in PyMOL (The PyMOL Molecular Graphics System, Version 1.8.4.0 Schrödinger, LLC).

## Results

### Purification of CII toxin

Con-ikot-ikot (CII) toxin was purified as described in (Chen et al., 2014). Each individual purification was started from 12 L of E. *coli* cultures. Despite large starting volumes, final toxin yield was modest, similar to previous reports (Chen et al., 2014): of three purifications, two resulted in ~100 μg and one in ~40 μg of CII toxin. The final product migrated as a dimer on a non-reducing SDS-PAGE gel (Fig. 2A-B). The CII dimer is ~20 kDa in size, but migrates like a larger protein on SDS-PAGE gels.

The activity of each batch of the purified toxin was tested in macroscopic recordings, either by perfusing AMPA receptors in an outside-out patch with 500 nM toxin and 10 mM Glu or by incubating HEK cells expressing AMPA receptors in a saturating concentration of the toxin (≥ 350 nM) (Fig. 2C). AMPA receptor desensitisation block was used as a measure of CII toxin activity, determined as the ratio of the steady-state current over the peak current. In a complete block of desensitization, the AMPA receptor current was sustained as long as glutamate was present, resulting in a square current response (Fig. 2C, right).

To determine the apparent binding affinity of the toxin, its ability to block AMPA receptor desensitization was tested at a range of different concentrations (0.3 – 300 nM) following a long incubation (~12 h). The resulting dose-response curve was fit with a Hill equation (see Materials and Methods) giving *EC*_50_ of 5 nM ± 2 nM and Hill slope of 1 ± 0.3 (at 95% confidence interval, Fig. 2D).

### Binding and unbinding of Con-ikot-ikot toxin to individual AMPA receptors

We next sought to observe binding and unbinding of CII toxin to individual AMPA receptors. To observe CII binding, AMPA receptors in a patch were perfused with 10 mM glutamate and 500 nM CII. In patches containing more than one AMPA receptor, binding of CII to only one receptor was regularly seen because the low CII concentration led to a slow association rate (Fig. 3A). Low toxin concentration also meant that patches must be perfused for minutes before a binding event was observed. Toxin binding was evident as a sudden change from a quiescent, desensitized state during the 10 mM glutamate pulse to frequent, brief openings and closures, without long, closed periods (Fig. 3A). Thus, the binding of toxin prevented receptors from desensitizing, although occasionally, longer closures of >23 ms (the longest shut time within a burst) were still present (Fig. 3B).

Once the toxin was bound (either via perfusion of CII or by incubation of the receptors in a saturating CII concentration), the patch was perfused in a solution containing 10 mM glutamate without any toxin to observe unbinding. An unbinding event was noted when AMPA receptor activity changed from frequent and brief openings and closures to long quiescent periods with rare bursts of activity (Fig. 3B). In agreement with the toxin’s high affinity, unbinding of toxin took 10.4 ± 5.5 min (*n* = 5) (Fig. 3C).

After establishing that the binding of toxin to AMPA receptors blocks desensitization, while at the same time resulting in short, but frequent closures, we next investigated how this behaviour compares to other desensitization blockers.

### Comparison of Con-ikot-ikot toxin to other desensitization blockers

Upon establishing that purified toxin is active and that it blocks AMPA receptor desensitization at single-channel level, we next sought to compare CII toxin to two other desensitization blockers: cyclothiazide (CTZ) and (R, R)-2b. CTZ is the most widely used AMPA receptor desensitization blocker (Partin et al., 1994). Two molecules of CTZ bind within one AMPA receptor LBD dimer, holding it together and preventing desensitization (Sun et al., 2002). At single-channel level, CTZ has been reported to increase open probability and burst duration, as well as decrease occupancy of the lowest conductance level and increase occupancy of higher conductance levels for both, GluA1 and GluA4 homomers (Fucile et al., 2006; Zhang et al., 2017). (R, R)-2b targets the same binding sites as CTZ, spanning two sites in a dimer with a double-headed arrangement, but is less characterised and used than CTZ (Kaae et al., 2007). A single molecule of (R, R)-2b binds per one AMPA receptor LBD dimer with *EC*_50_ of 0.44 μM, an order of magnitude tighter binding than the *EC*_50_ of CTZ (5 μM). We tested (R, R)-2b here as it has been used in several AMPA receptor structures with (Chen et al., 2014) and without toxin (Dürr et al., 2014).

To compare the effects of CTZ, (R, R)-2b and Con-ikot-ikot toxin on AMPA receptor single-channel activity, we generated all-point-amplitude histograms from recordings of individual GluA2 receptors made in each of the three conditions (Fig. 4). The histograms were made from parts of the recordings where the receptor was perfused with glutamate and the corresponding modulator, so any closed periods (the baseline around 0 pA) are a consequence of the receptor inactivity during agonist application (see also Materials and Methods). The histograms show there is heterogeneity in the channel activity between patches for each positive modulator. For example, in CTZ, 2 out of 8 patches showed pronounced peaks in the histogram at 0 pA indicating the receptor spent considerable amounts of time in non-conducting states. Some patches in CTZ showed receptor being open most of the time, rarely reaching the baseline level of 0 pA. The majority of patches in CTZ, however, showed a sustained receptor activity at lower conductance levels, rarely opening to >2 pA at –80 mV. In sharp contrast, modulation by (R, R)-2b enabled the GluA2 receptor to open consistently to amplitudes over 2 pA at –80 mV. In this context, Con-ikot-ikot toxin was revealed as the poorest promoter of the active state of AMPA receptors. When complexed with the CII toxin, the receptor frequently visited non-conductive states in the presence of 10 mM glutamate in all patches tested. At the same time, visits to open amplitudes above 2 pA at −80 mV were less frequently observed than in CTZ and especially in (R, R)-2b.

**Figure 4.**
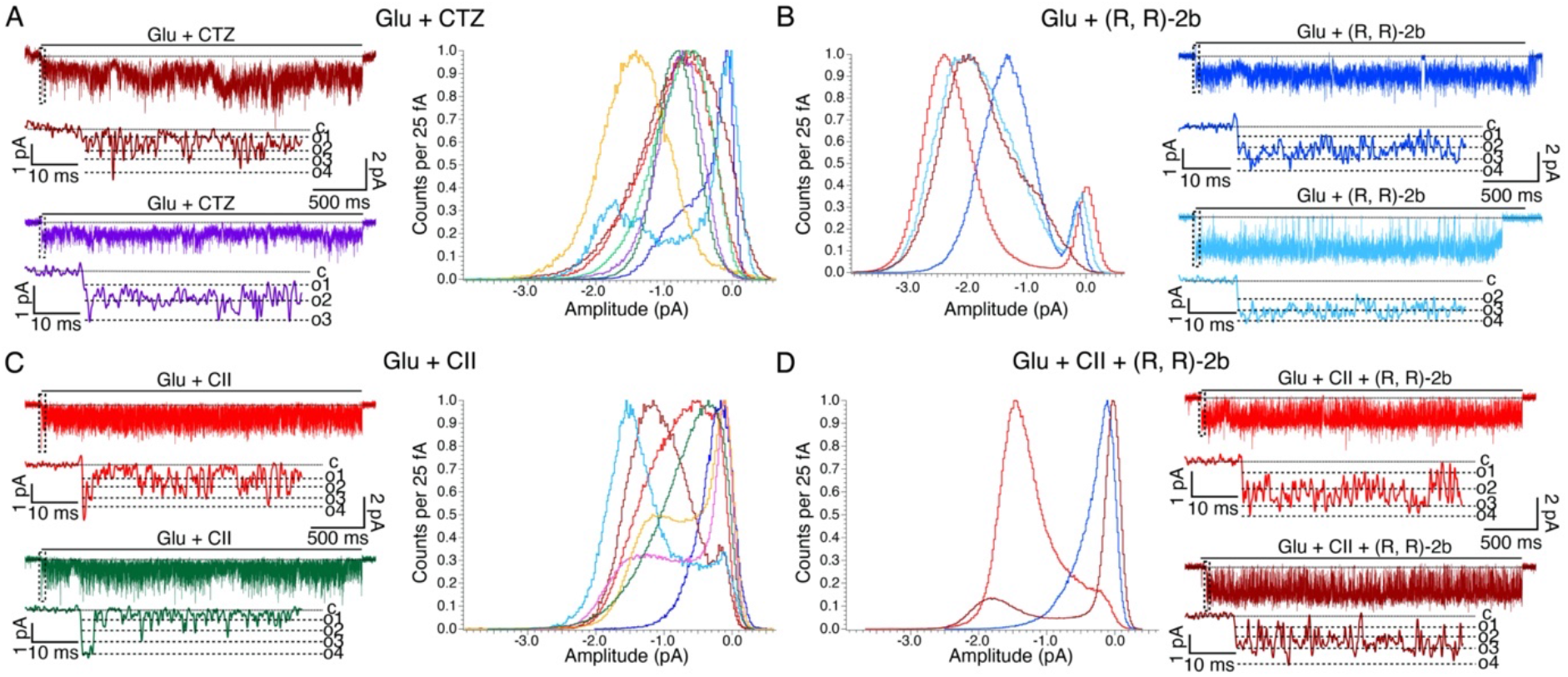
Comparison of Con-ikot-ikot toxin with AMPA receptor desensitization blockers CTZ and (R, R)-2b. (A) All-point-amplitude histograms of single-channel recordings of GluA2 wild-type receptors perfused in glutamate and positive modulator cyclothiazide (CTZ), *n*_CTZ_ = 8 (~46 sec of recording in total). A histogram was made for each patch, using only the parts of the recording where the patch was perfused in glutamate (Glu) and CTZ. Histograms were normalised to the maximum value. Representative single-channel currents from colour-coded patches are drawn next to each histogram. Dashed box indicates the part of the trace shown at greater magnification below. Dotted lines are baseline (closed level (c) in zoom), and dashed lines are open levels in each patch (O1-O4). All openings are downwards, traces were obtained at ~ −80 mV and low pass filtered at 1 kHz for presentation. (B-D) Same as in (A), but with a different modulator: (R, R)-2b (*n*_RR_ = 4; ~81 sec of recording in total), Con-ikot-ikot toxin (*n*_CII_ = 7; ~77 sec of recording in total) and (R, R)-2b + CII (*n*_RR+CII_ = 3; ~48 sec of recording in total), respectively.

The toxin, binding a single site on the top of the LBDs, was the least efficient modulator in promoting higher conductance states and allowed frequent closures. Both CTZ and (R, R)-2b bind within the intra-dimer LBD interface. In principle, toxin and (R, R)-2b should not occlude each other’s binding, as seen in a complex captured in crystal structures, together with a partial agonist (Chen et al., 2014). In order to test whether immobilisation of the LBD layer by toxin leads to an overall decrease in activity of AMPA receptors, even in the presence of other desensitization blockers, we next sought to record activity of single AMPA receptors in the presence of both toxin and (R, R)-2b.

### (R, R)-2b cannot overcome the low AMPA receptor activity induced by toxin binding

In order to investigate whether immobilisation of the LBD layer by toxin is essential for its ability to reduce AMPA receptor activity, we recorded currents through single AMPA receptors treated with both, toxin and (R, R)-2b. If locking the two LBD dimers with respect to each other by the toxin is indeed preventing AMPA receptors from stably occupying the most open levels, then the presence of the toxin should have a negative impact on AMPA receptor activity even in the presence of a potent positive modulator, such as (R, R)-2b. The challenging aspect of these recordings was to know that both modulators were bound to the receptor. In majority of the recordings, the coverslip with HEK293 cells expressing wild-type GluA2 receptors was incubated in a solution containing saturating toxin concentration. Almost all of patches from such cells contained receptors bound by the toxin, as evident from the characteristic single-channel activity induced by bound toxin (Fig. 3). Patches with bound toxin were then perfused in saturating glutamate (10 mM) and (R, R)-2b (10 μM) for at least 10 s. Due to high affinity and saturating concentration of (R, R)-2b, it binds to AMPA receptors almost instantaneously, so this incubation should be sufficient for (R, R)-2b to saturate a few AMPA receptors in a patch. The reverse experiment, where (R, R)-2b was first allowed to bind to the receptors and subsequently toxin was added could not be performed satisfactorily. Patch recordings are unreliable and perfusing patches that did not give single channel recordings consumed prohibitive amounts of the toxin preparation. Crystal structures of AMPA receptors in complex with the toxin, (R, R)-2b and a partial agonist, where the receptors were first incubated in Con-ikot-ikot toxin (at 1:1.5 moral ratio of receptor to toxin, respectively) and then (R, R)-2b suggest (R, R)-2b can access its binding sites in toxin-bound AMPA receptors (Chen et al., 2014).

All-point-amplitude histograms for receptors treated with toxin and (R, R)-2b show more similarity to toxin than (R, R)-2b-bound receptors as the receptor rarely reaches amplitudes over 2 pA at –80 mV (Fig. 4D). In all-point-amplitude histograms, the area under each peak is proportional to the time spent at the given amplitude, but there is no indication of the duration of individual amplitudes. In other words, the histograms do not distinguish between frequent and short openings vs. rare and long openings to a specific amplitude. To be able to better compare whether the presence of (R, R)-2b changes activity of toxin-bound AMPA receptors, we idealised the single-channel records using the threshold crossing method (see Materials and Methods), using multiple thresholds between user-defined open levels (Fig. 5D). Four distinct open levels were identified for receptors bound by toxin (O1: −0.59 pA ± 0.01; O2: - 1.20 pA ± 0.02; O3: −1.75 pA ± 0.04; O4: −2.31 pA ± 0.08; *n* = 4 patches), bound by (R, R)-2b (O1: −0.58 pA ± 0.00; O2: −1.19 pA ± 0.00; O3: −1.77 pA ± 0.04; O4: −2.5 pA ± 0.08; *n* = 3) or exposed to both CII and (R, R)-2b (O1: −0.59 pA ± 0.00; O2: −1.20 pA ± 0.00; O3: −1.78 pA ± 0.01; O4: −2.37 pA ± 0.01; *n* = 3). These data are quite similar to those reported previously for GluA2 with CTZ bound (Prieto & Wollmuth, 2010). With only toxin bound to its LBDs, the receptor spent the least fraction of time (3.9% ± 1.7) at the highest amplitude level, level 4, indicating rare and short-lived visits to the highest amplitude state. The receptor spent approximately equal fractions of time across the closed state and remaining 3 open sublevels (C: 26 ± 7%; O1: 25 ± 8%, O2: 26 ± 5%; O3: 19 ± 7%; Fig. 5A, bottom). If (R, R)-2b was used as desensitization blocker, the occupancy profile of the receptor changed, with biggest differences at the closed, O1 and most open level (O4). The receptor now spent almost 10-fold less time in a closed state, compared to the CII-bound receptor (CII: 26 ± 7%, (R, R)-2b: 3 ± 2.5%) and occupancy of the lowest open level (O1) was ~3 times lower (CII: 25 ± 8%, (R, R)-2b: 9 ± 6%), whereas the fraction of time spent at level 4 (O4) was ~10 times greater (CII: 4 ± 2%, (R, R)-2b: 40 ± 19%) (Fig. 5B, bottom). For AMPA receptors exposed to both toxin and (R, R)-2b, the level occupancy is best described as intermediate between toxin and (R, R)-2b profiles, with the receptor spending 28 ± 22% of the time in the closed state, similar to when only toxin is bound, and 9 ± 9% of the time at most open level, a value higher than when only toxin is bound and lower than when only (R, R)-2b is bound (4% and 40% respectively see above) (Fig. 5C). The occupancy of the intermediate open levels, O2-O3, was not much different in toxin or (R, R)-2b, in each case, each level being occupied for 20-25% of the time (Fig. 5A-C).

**Figure 5.**
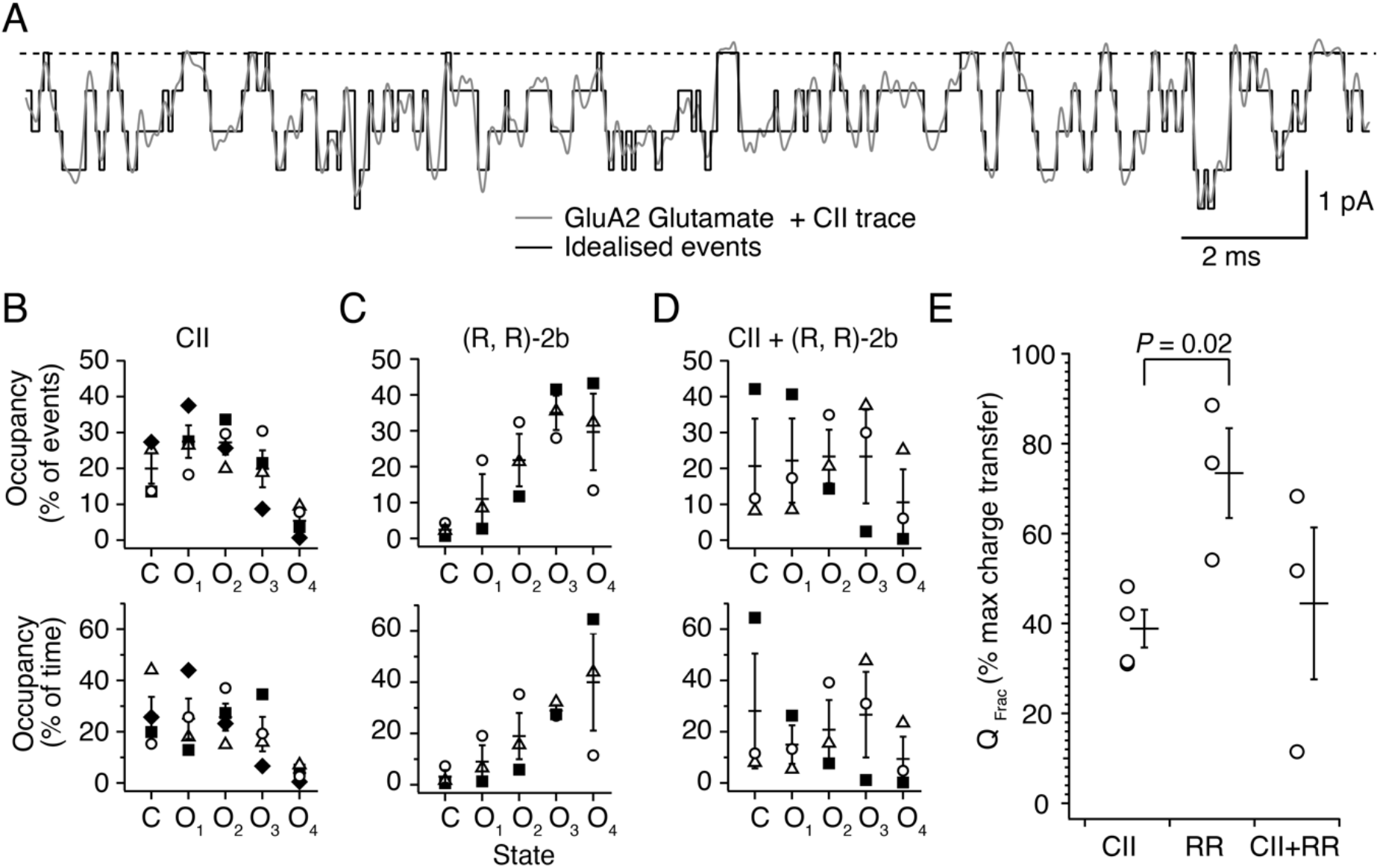
Occupancy of closed and open levels for GluA2 bound by modulators of desensitization. (A) An example of the idealisation from ASCAM (black), overlaid on the current trace (grey; filtered with 1 kHz LPF for presentation) obtained in glutamate (Glu) and toxin (CII). Dashed line indicates baseline. (B-D) Four different open levels were detected in (B) toxin alone, (C) (R, R)-2b and (D) toxin + (R, R)-2b conditions. Occupancy of each level (number of visits normalized to the total number of events, upper row) and occupancy of each level as a fraction of the total time (bottom row) are shown. The presence of both modulators produced an intermediate, heterogeneous effect. Different symbols in each condition represent different patches. (E) The percentage of maximum charge transfer was determined as the ratio of the transferred charge to the charge that would be transferred if the channel were open continuously to the maximum open amplitude; CII 38 ± 4% (*n* = 4), (R, R)-2b 73 ± 10% (*n* = 3), and toxin and (R, R)-2b 44 ± 17% (*n* = 3). Horizontal bars are mean values and bars represent standard error of the mean. Probability of no difference is from Student’s two-tailed *t*-test.

Single channel activity is often expressed as open probability – the fraction of time in open states compared to the overall time. Because GluA2 shows 4 evenly-spaced sublevels when desensitization is blocked, this measure (which weights all openings equally independent of their amplitude) is somewhat inadequate. To assess “openness” (or activity) of the channel in each condition, we determined the fraction of maximum charge (*Q*_Frac_) passed by the channel as described in Materials and Methods. In the presence of toxin, the channel transferred 38 ± 4% of the maximum charge (*n* = 4). This value almost doubled in (R, R)-2b (73 ± 10%, *n* = 3), whereas the presence of both modulators resulted in an intermediate value (44 ± 17%, *n* = 3) with a wider range of values (Fig. 5E). The probability of no difference between CII and (R, R)-2b was 0.02 (Student’s two-tailed *t*-test).

To understand the effects on receptor activation more clearly, we next generated dwell time histograms for each condition (Fig. 6). All histograms were fitted with two exponential components, the values of which are summarised in Table 1. The durations of different open levels and closed states are comparable across the 3 conditions, and the presence of toxin or (R, R)-2b or both modulators did not change them. Thus, the modulators do not change the amplitudes and durations of visits to open and closed states, but exert their effect via shifting frequency of visits to the closed state and the 4^th^ (largest amplitude) open level.

**Figure 6.**
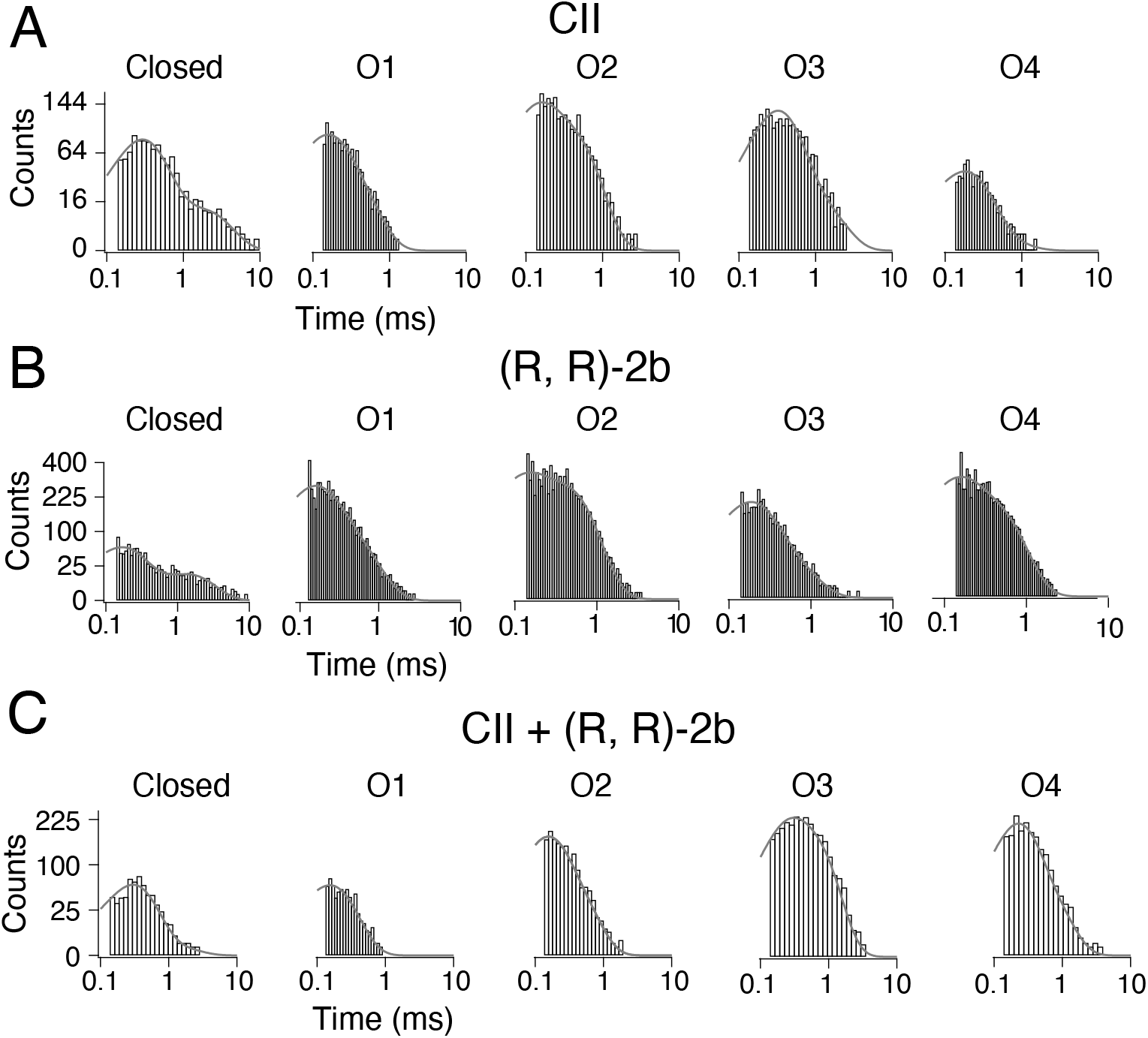
Dwell times of AMPA receptors bound by Con-ikot-ikot toxin in the presence and absence of (R, R)-2b. (A-C) Dwell time histograms were generated for the closed and each of the open levels, for (A) toxin, (B) (R, R)-2b and (C) toxin and (R, R)-2b bound to the receptor. Shown are histograms from one representative patch for each condition. Each histogram was plotted on a log scale and fit with two exponential components (grey curves). Time constants and amplitudes are given in Table 1. O1-4 indicate open levels 1-4.

**Table 1.** Time constants from exponential fits to dwell time histograms. Single-channel recordings of GluA2 wild-type receptors were obtained in glutamate and the following positive allosteric modulators: Con-ikot-ikot toxin (CII), (R, R)-2b or the combination of the two. Dwell times of closed and open levels (O_1-4_) were obtained by idealising single-channel traces and each dwell time histogram shown in Fig. 6 was fit with 2 exponential components (τ_1_ and τ_2_); “A_1_” indicates amplitude of the first component (τ_1_) expressed in % and *n* number of patches.

| Dwell times ($\tau$) in $\mu\text{s}$ | | CII<br>$n = 4$ | (R, R)-2b<br>$n = 3$ | CII + (R, R)-2b<br>$n = 3$ |
| --- | --- | --- | --- | --- |
| Closed | $\tau_1$ | $250 \pm 30$ | $164 \pm 12$ | $279 \pm 5$ |
| | $\tau_2$ | $1700 \pm 400$ | $900 \pm 300$ | $1000 \pm 250$ |
| | $A_1(\%)$ | $73 \pm 4$ | $73 \pm 6$ | $75 \pm 16$ |
| $O_1$ | $\tau_1$ | $118 \pm 13$ | $129 \pm 5$ | $109 \pm 20$ |
| | $\tau_2$ | $400 \pm 60$ | $360 \pm 80$ | $300 \pm 70$ |
| | $A_1(\%)$ | $48 \pm 6$ | $61 \pm 5$ | $47 \pm 12$ |
| $O_2$ | $\tau_1$ | $150 \pm 30$ | $96 \pm 16$ | $129 \pm 11$ |
| | $\tau_2$ | $500 \pm 90$ | $360 \pm 50$ | $400 \pm 60$ |
| | $A_1(\%)$ | $53 \pm 13$ | $52 \pm 9$ | $54 \pm 10$ |
| $O_3$ | $\tau_1$ | $160 \pm 40$ | $74 \pm 16$ | $133 \pm 13$ |
| | $\tau_2$ | $480 \pm 200$ | $390 \pm 20$ | $420 \pm 130$ |
| | $A_1(\%)$ | $40 \pm 15$ | $51 \pm 8$ | $41 \pm 5$ |
| $O_4$ | $\tau_1$ | $174 \pm 17$ | $197 \pm 36$ | $173 \pm 3$ |
| | $\tau_2$ | $400 \pm 100$ | $900 \pm 300$ | $450 \pm 150$ |
| | $A_1(\%)$ | $83 \pm 4$ | $55 \pm 6$ | $66 \pm 16$ |

## Discussion

The rich palette of positive modulators of AMPA receptors spans a range of chemical complexity from ions (Dawe et al., 2016; Partin et al., 1996) to small molecules (Vyklicky et al., 1991; Yamada & Tang, 1993) with Con-ikot-ikot toxin being the largest known. All act on the LBD layer, and have been largely characterised by macroscopic measurements of AMPA receptor currents. In such measurements, modulator activity is defined only relative to the original response, or other modulators. Many AMPA receptor positive modulators are highly lipophilic (e.g. cyclothiazide), making washout difficult and meaning relative measurements are confounded by loss of activity. Single channel measurements, in contrast, allow an absolute definition of activity, and also give insight into mechanism. Without auxiliary proteins, GluA2 has an activity (expressed as steady-state current) of about 3% in saturating glutamate (Carbone & Plested, 2012). As a measure of activity that also takes into account amplitudes occupied by the open channel, we determined the fraction of maximum charge (*Q*_Frac_) passed by the channel in single-channel records. Our measurements show that the addition of *in vitro* purified CII toxin increases AMPA receptor activity, and that the increase is submaximal (only about 13-fold, to 38% of the maximum), less than that produced by other small molecule blockers of desensitization (24-fold increase to about 73% of maximal activity for (R, R)-2b). When both, toxin and (R, R)-2b are bound to the receptor, the increase in activity is about 15-fold, to 44%. This finding explains, at least in part, why toxin bound crystal structures with partial agonists and (R, R)-2b had closed ion channel pores. In these conditions, the energy landscape remains tilted in the direction of inactive states.

Although the toxin blocked desensitization, prolonged shut periods characteristic of desensitized receptors were present. These closures, longer than the cutoff value of 23 ms, are most likely desensitization events, although partial toxin unbinding (from some, but not all subunits), followed by reassociation, cannot be excluded. This residual desensitization is not unique to toxin and was reported for AMPA receptors in complex with CTZ, and those carrying single-point mutations that block desensitization (LY and Lurcher mutations) (Zhang et al., 2017). Even though the main mechanism of desensitization in AMPA receptors proceeds via rupture of the LBD dimers (Sun et al., 2002), desensitization from the LBD-TMD linkers has been reported previously (Yelshansky et al., 2004).

We observed 4 different open levels for GluA2, either with toxin and/or (R, R)-2b bound, just like freely-desensitizing AMPA receptors (Zhang et al., 2008). Receptors in complex with CII toxin shut frequently, gave a flat occupancy distribution across the 3 smallest amplitude sublevels, and rarely visited the maximum open level. Comparison of the toxin with other desensitization blockers, CTZ and the more potent (R, R)-2b (Kaae et al., 2007), showed that frequent closures and a paucity of full-amplitude openings are characteristic for the toxin. When CII toxin is bound, receptors spend about 10-fold more time in the closed state and about 10-fold less time in the 4^th^ open level compared to (R, R)-2b-bound AMPA receptors. Different occupancies of the closed and 4^th^ open level are the main mechanistic difference between the toxin and (R, R)-2b. A similar effect has also been reported for CTZ when compared to freely desensitizing receptors (Fucile et al., 2006; Zhang et al., 2017). There was no difference in open channel amplitudes, nor in the time constants describing the closed and open levels. This indicates that (R, R)-2b is a better stabiliser of the full open state because it allows the receptors to reach the 4^th^ open level more frequently, primarily at the expense of the closed state. Concomitantly, with the toxin bound, receptors struggled to reach the fully-activated state as often. In our recordings, exposure to both toxin and (R, R)-2b gave an intermediate effect, indicating that both toxin and (R, R)-2b are bound to the receptor, and that some within-dimer effects that occur in CII-bound receptors can be reverted by (R, R)-2b.

Given that sublevel occupancy is related to agonist binding to subunits that activate independently (Rosenmund et al., 1998) and we worked in saturating glutamate (10 mM), this presents a conundrum about how CII works. The toxin binds at the opposite side of the LBD layer to the LBD-TMD linkers, and it seems unlikely that the toxin prevents agonist binding (Chen et al., 2014). Therefore, LBD inter-dimer angle and lateral displacement remain as mechanisms that permit the toxin to block desensitization, and at the same time fail to promote occupancy of the highest conductance state (Fig. 7). Con-ikot-ikot is a homodimer, with a flattened V-shape. When bound, the toxin immobilises the two LBD dimers at a fixed angle of ~35°, similar to what is observed in apo receptors (Chen et al., 2014; Dürr et al., 2014). In the absence of toxin, the two LBD dimers relax, adopting a ~55° inter-dimer angle. Previous work showed that cross-links formed between the LBD dimers in AMPA receptors, which inhibit lateral movements, lead to a decrease in receptor activity (Baranovic & Plested, 2018; Baranovic et al., 2016), irrespective of the nature (disulphide or zinc bridges or flexible bifunctional cross-linkers) or position of the bridge. A similar observation has been reported for closely related kainate receptors where disulphide bridges across the LBD dimers locked receptors in a semi-active conformation (Daniels et al., 2013). In contrast, a disulphide bridge at the LBD dimer interface locks NMDA receptors in a super-active state (Esmenjaud et al., 2019), but this must be considered in the context of standing inhibition by the ATD layer, which appears absent in AMPA receptors. The opposing effects of CII toxin to block desensitization and also inhibit activity, whilst still allowing full opening, suggest that, in the absence auxiliary subunits, AMPA receptors sample a set of conformations when the channel pore is fully conducting. LBD layer dynamics, and a conformation distinct from that stabilized by the toxin, are key for high activity.

**Figure 7.**
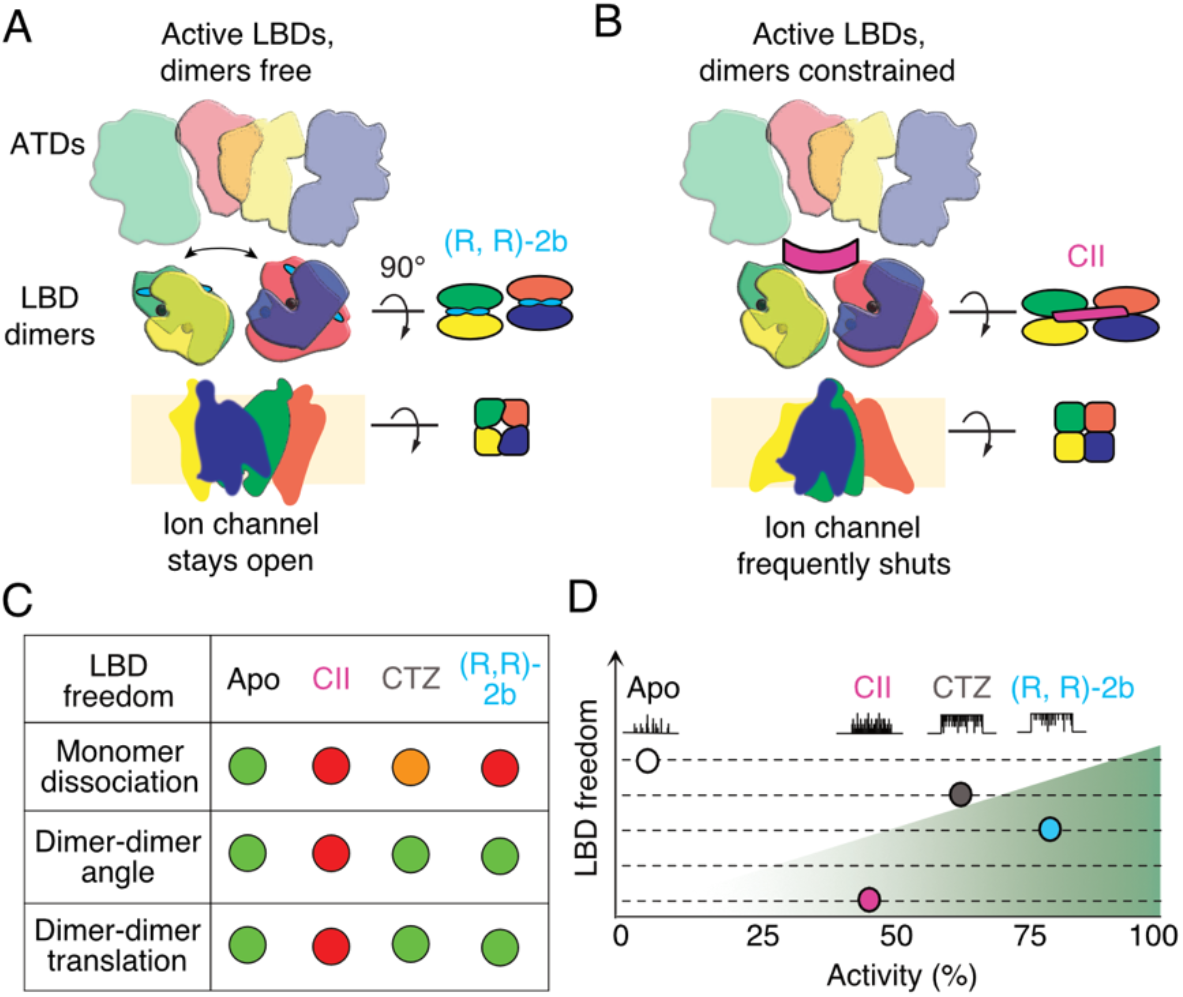
Conformational freedom of LBD layer dictates activity of AMPA receptors. (A) AMPA receptor subunits are colour-coded, with green and yellow forming one ligand binding domain (LBD) dimer, and blue and red another. AMPA receptors freely desensitize as long as the dimers are allowed to disassociate, resulting in very low activity of wild-type receptors. Fixing the two monomers within each dimer, by CTZ or (R, R)-2b (cyan, 2 bars), blocks AMPA receptor desensitization and increases activity. (B) Preventing lateral movements and/or rotations of the LBD dimers, through binding of Con-ikot-ikot toxin (CII, magenta) lowers the activity of the ion channel. (C) Summary of degrees of freedom of the LBD layer. (R, R)-2b binds with higher apparent affinity to LBD dimers than CTZ. (D) Fictive single channel currents indicate increasing activity over the four conditions. (R, R)-2b supports the highest activity because it holds LBD dimers tightly together whilst allowing their free rotation and lateral translation to the optimal position.

Gating modes are observed for AMPA receptors (Prieto & Wollmuth, 2010) and for receptors in complex with auxiliary proteins (Zhang et al., 2014). We saw clear heterogeneity in our recordings, with non-desensitising channels exhibiting qualitatively 2 modes of gating: one where openings were mostly to the higher amplitudes and a less frequent one, where openings were mainly to lower conductance levels. In case of CII recordings, the low conductance mode was observed in 2/7 patches, and in both patches, it was the dominant mode, accounting for >50% of the gating time. Although we do not know what the origins of these different modes are, we do know they are not specific for the toxin, as we have also observed them with the other modulators (CTZ and (R, R)-2b). Here we analysed exclusively high activity modes, because they resemble recordings in previous reports much better than the low activity modes generally do. High and low open probability gating modes have been previously reported for AMPA receptors complexed with CTZ, where they were related to voltage, but in saturating (5 mM) glutamate, only the high gating mode was reported (Prieto & Wollmuth, 2010). In contrast, we observed both modes in saturating (10 mM) glutamate. A possible explanation for this could be that the gating modes change more slowly at high concentration, and in previous reports, the recordings were too brief to sample them (in the range of seconds, (Prieto & Wollmuth, 2010). Longer recordings might increase the probability of observing rare channel behaviours, such as low open probability gating mode in saturating glutamate concentration (Zhang et al., 2014). During long-lived recordings, patches might undergo local invaginations or other spatial rearrangements which might decrease effective local agonist concentration and lead to occurrence of low open probability gating mode in saturating glutamate (Suchyna et al., 2009).

Toxins have been used extensively as tools to study ion channel function (Kalia et al., 2015) and our work shows that the very high affinity of the toxin should allow it to be used for experimental applications similar to antibodies, but without the associated bulk. Con-ikot-ikot is specific for AMPA receptors – it can bind to any of its subunit types (GluA1-4), but does not have an effect on the closely-related kainate (GluK2) or NMDA receptors (GluN1/GluN2A) or on GABA-A receptors (Walker et al., 2009). The toxin also blocks desensitization of native, likely heteromeric, AMPA receptors, which is unsurprising given that GluA2 residues interacting with the toxin are conserved across all four subunits (Chen et al., 2014). The observation that the toxin actually inhibits the maximum activity provides insight into how the toxin works *in vivo.* The inhibition of desensitization implies that CII exploits a large extrasynaptic receptor pool, and that a key physiological role of desensitization is to mask the potentially deadly capacity of this receptor pool for generating depolarizing current. The initial report about CII (Walker et al., 2009) estimated the *EC*_50_ to be 67 nM. This value was an underestimation, likely due to the lack of time to equilibrate in the oocyte recording system. The high affinity we determined with overnight incubation (5 nM) indicates that the toxin should tolerate mutations and derivatizations that would diversify its functionality whilst still binding well to AMPA receptors. Congruent with its high affinity, the toxin stays bound to GluA2 homomeric receptors for an average of 10 minutes. Due to its small size and unique binding site, it leaves the overall dimensions of the receptor unchanged and extracellular domains free to interact with other synaptic proteins, unlike antibodies or even Fab fragments labelling the ATD (Giannone et al., 2010; Zhao et al., 2019). Although further characterisation of the toxin is needed, such as possible effects of different auxiliary proteins, these properties suggest toxin has potential as a tool for investigating native AMPA receptors.

## Author Contributions

JB purified the toxin, recorded macroscopic and single channel currents, and analysed data;

SB recorded macroscopic and single channel currents and analysed data, SM and MC purified the toxin, AJRP and JB wrote the manuscript with input from all authors

## Acknowledgements

We thank Eric Gouaux for sharing the toxin purification protocol, plasmid and (R, R)-2b, Lei Chen for help with toxin purification, Marcus Wietstruk for molecular biology and Remigius Lape for comments on the manuscript.

